# Developmental remodeling of ping-pong piRNA amplification in the vertebrate female germline

**DOI:** 10.64898/2026.09.10.750710

**Authors:** Mohammad H. Ghazimoradi, Ali Nemati, Cheng Ma, Lijiang Tang, Sofia Corazza, Sara Yousefi Teameh, Amir Fallahshahroudi

## Abstract

The piRNA pathway silences transposable elements (TEs) in the germline, and the ping-pong amplification cycle is the hallmark of this defense. In the male germline, ping-pong is most active during a meiotic window of spermatogenesis, yet its developmental profile in the vertebrate female germline remains less well explored. Most profiling has used adult ovary and mature oocytes, stages at which piRNA pathway components are reported to be low. To address this, we generated matched strand-specific RNA-seq and small RNA-seq from pre-meiotic (E10.5) and meiotic entry (E16.5) chicken ovary, used published single-cell data to track germ-cell composition across the same window, and extended the analysis to the mature chicken ovary and to zebrafish across developmental stages. Ping-pong amplification increases at meiotic entry compared to the pre-meiotic stage across TE classes. In the mature ovary, the signature weakens, and the remaining ping-pong pairs are preferentially associated with LTR/ERV retroelements. We show that activation of a meiotic entry transcriptional program in an *in vitro* chicken primordial germ cell model increases the fraction of piRNA-sized reads with a partner exhibiting a 10-nt 5ʹ overlap and increases the 1U signature of piRNA-sized reads, consistent with meiotic priming promoting piRNA biogenesis. The zebrafish ovary shows a similar meiosis-associated amplification and preferential targeting of LTR/ERV retroelements at maturity, while carrying roughly 5.7-fold more TE sequences. Similar patterns in two lineages that diverged approximately 430 million years ago suggest that germline development shapes both the timing of ping-pong amplification and the TE classes preferentially associated with it.

## Introduction

Fertility and species continuity depend on maintaining germline genomic integrity. This integrity is continually challenged by transposable elements (TEs), mobile genetic elements that can disrupt gene function and compromise genome stability (1). To counteract this threat, animals have evolved the PIWI- interacting RNA (piRNA) pathway, a conserved small RNA-based surveillance system that silences TEs through two complementary strategies (2). In the nucleus, piRNA-guided PIWI proteins direct transcriptional gene silencing by recruiting chromatin-modifying complexes that establish repressive histone marks, particularly H3K9me3, and in some lineages catalyze de novo DNA methylation at TE loci(3). In the cytoplasm, the pathway enforces post-transcriptional silencing through ping-pong amplification, a reciprocal mechanism in which PIWI-piRNA complexes cleave TE transcripts and complementary precursors to generate secondary piRNAs. An antisense piRNA, often bearing a 5ʹ uridine bias (1U), is loaded into a PIWI protein and directs endonucleolytic cleavage of a complementary sense TE transcript. The cleavage product is processed into a sense secondary piRNA, often bearing an adenine at position 10 (10A), and loaded into a PIWI protein (4). The resulting PIWI-piRNA complex can then cleave antisense precursors, regenerating antisense piRNAs and sustaining the loop. The hallmark of this active cycle is a precise 10- nucleotide 5ʹ overlap between complementary sense and antisense piRNA pairs, which is widely used as the canonical sequence signature of ping-pong amplification (5).

In *Drosophila melanogaster*, where the ping-pong cycle was first described, Aubergine and Argonaute3 drive cytoplasmic amplification in ovarian nurse cells, while nuclear Piwi enforces co-transcriptional silencing at TE loci (6–8). In mice, an earlier, pre-pachytene population of piRNAs, produced largely in mitotically arrested fetal prospermatogonia before meiotic entry, is enriched for TE-targeting sequences and drives ping-pong amplification and de novo TE methylation; a later pachytene population is largely, though not entirely, depleted of TE targets (9, 10). This stage resolution matters; it establishes that, in the male germline, TE-directed ping-pong is not constitutive but is engaged during a defined developmental window, yet this model is not well studied in females. It has been shown that PIWI proteins and piRNAs are present in rodent and human oocytes, where they contribute to TE suppression through predominantly post-transcriptional mechanisms (11, 12). In mice, loss of PIWI proteins causes male sterility, but females remain fertile (13), while in the golden hamster, loss of Mov10l1 or Piwil1 causes sterility in both sexes (14). The mouse phenotype should therefore not be generalised to other mammals.

In birds, the evidence for ping-pong amplification comes mainly from the male germline (15, 16). In chicken (*Gallus gallus*), two PIWI homologs, PIWIL1 (CIWI) and PIWIL2 (CILI), mediate a robust ping-pong cycle in the testis that preferentially targets transcriptionally active TE families, particularly endogenous retroviruses (ERVs) and CR1 non-LTR retrotransposons. Stage-resolved profiling of the first wave of spermatogenesis has since shown that this testis program is itself developmentally gated, with the piRNA burst emerging at the pachytene stage (16, 17). Profiling of the adult chicken ovary reported that PIWI genes and piRNA precursors are essentially undetectable there, leading to the proposal that the avian piRNA pathway is a sexually dimorphic, testis-restricted feature (17, 18). This strict dimorphism model is not universal in amniotes. A survey of piRNA-pathway genes found conserved expression of PIWIL2, PIWIL4, and MAEL in oocytes and ovarian supporting cells of human, mouse, and platypus, though the same study found no PIWIL1 in either mouse or chicken ovary, consistent with the chicken-specific negative finding above (15). In the chicken, female primordial germ cells (PGCs) arrive at the gonad around embryonic day 3-5 (E3-5) (19, 20), undergo mitotic expansion through E10.5, activate the meiotic program from E12.5, and enter meiosis around E16 (SPO11^+^). Because these adult-ovary studies sampled follicle-dominated tissue, a germline piRNA signal could be diluted by the somatic compartment rather than genuinely absent. Consistent with this, knockdown of PIWIL1 and PIWIL2 in cultured female PGCs derepresses CR1 retrotransposons and induces DNA double-strand breaks, establishing a protective role for these components (21), and more recent work detecting piRNAs in female embryonic PGCs and oocytes, together with PIWIL1 and PIWIL2 expression throughout female PGC stages, is consistent with a pathway that is developmentally transient rather than constitutively undetectable (22, 23).

Here, we combined matched strand-specific RNA-seq and small RNA-seq with published single-cell RNA-seq to follow the female piRNA program through chicken ovary development, from mitotic (E10.5) to meiotic-entry (E16.5) and mature ovaries, and compared it with zebrafish at pre-meiotic, meiotic, and mature stages. We also asked whether inducing the meiotic program in cultured chicken PGCs changes piRNA biogenesis and ping-pong signatures.

## Results

### Meiotic entry couples piRNA-pathway induction to a transposon burst in the female chicken ovary

To define the transcriptomic changes accompanying the mitotic-to-meiotic transition, we performed differential expression analysis between E10.5 (mitotic) and E16.5 (meiotic entry) ovaries (DESeq2; n = 3). We identified 1,942 differentially expressed genes (1,364 upregulated, 578 downregulated; adjusted p < 0.05, |log_2_FC| ≥ 1; Table S1; Fig. 1A). Gene Ontology analysis of the upregulated set showed significant enrichment of reproductive process, sexual reproduction, and gametogenesis terms (Fig. 1B; adjusted p < 0.05). Consistent with meiotic entry, the E16.5 transcriptome showed a sharp downregulation of the pluripotency markers NANOG and SOX2, alongside a robust induction of the meiotic-initiation factors *STRA8* and *MEIOC* (Fig. 1C; adjusted p < 0.05). This meiotic switch was accompanied by induction of the piRNA pathway. *PIWIL1* and the associated factors *TDRD1*, *TDRD5*, *TDRD9*, *TDRD12*, *TDRKH*, *M1AP*, *MAEL*, and *MOV10L1* were all significantly upregulated at E16.5 (Fig. 1C; adjusted p < 0.05). We confirmed these changes by qRT-PCR in whole ovaries. Relative to E10.5, *STRA8* increased 6.2-fold, *SPO11* 7.7-fold, and *PIWIL1* 4.3-fold at E16.5, whereas *NANOG* was downregulated compared to E10.5 levels (Fig. 1D; two-tailed Student’s t-test, p < 0.05, n = 3).

**Figure 1.**
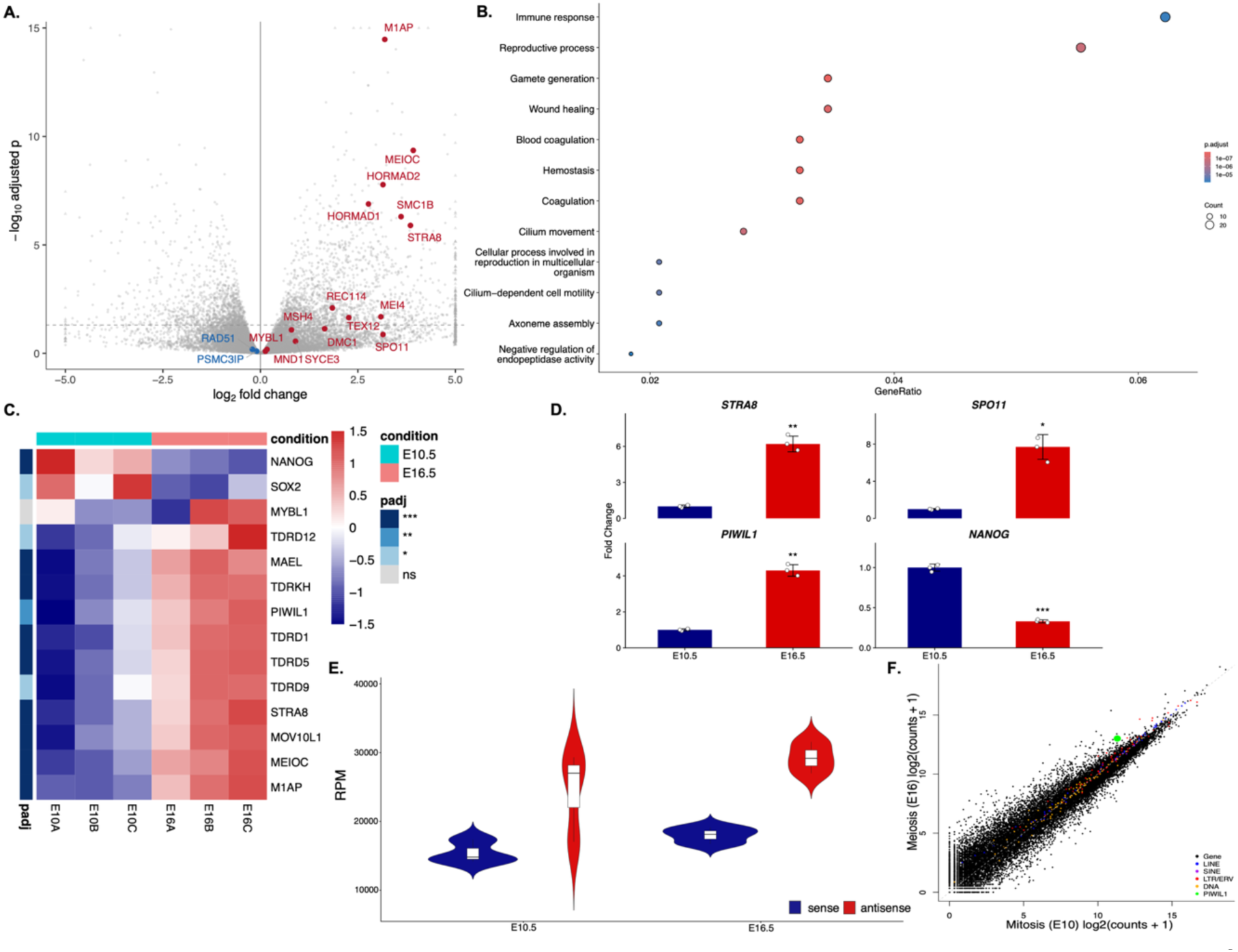
Meiotic entry is associated with upregulation of TEs and piRNA pathway genes in the female chicken ovary. **A.** Volcano plot of E10.5 (mitotic) versus E16.5 (meiotic entry) ovaries (DEGs; adjusted p < 0.05, |log_2_FC| ≥ 1) (bulk RNA-seq, n = 3 per stage, DESeq2). Meiosis-related genes upregulated at E16.5 are shown in red; downregulated genes in blue. **B.** Gene Ontology biological process enrichment bubble plot for DEGs upregulated at E16.5 (adjusted p < 0.05; Table S1). Bubble size indicates gene count; color indicates adjusted p-value (red = more significant; clusterProfiler, Benjamini-Hochberg). **C.** Heatmap of row-scaled normalized expression for piRNA pathway (*PIWIL1*, *TDRD1*, *TDRD5*, *TDRD9*, *TDRD12*, *TDRKH*, *M1AP*, *MYBL1*, *MAEL*, *MOV10L1*), meiotic (*STRA8*, *MEIOC*), and pluripotency (*NANOG*, *SOX2*) genes across individual replicates at E10.5 and E16.5. piRNA-pathway and meiotic genes are coordinately induced at E16.5, with reciprocal suppression of pluripotency markers. **D.** qRT-PCR validation of key markers (*STRA8*, *SPO11*, *PIWIL1*, *NANOG*) in whole ovaries at E10.5 (mitotic, blue) and E16.5 (meiotic, red). Expression is shown as fold change relative to E10.5 (mean ± SD, n = 3 biological replicates; *GAPDH* and *ACTB* as internal controls; two-tailed Student’s t-test: * p < 0.05, ** p < 0.01, *** p < 0.001). **E.** Violin plot of total TE-derived transcript abundance (RPM) from sense (Blue) and antisense (Red) strands at E10.5 (mitotic) and E16.5 (meiotic). Violins with boxplots showing median and interquartile range. Two-tailed Student’s t-test. Sense TE transcription increased significantly at E16.5 (p < 0.05), indicating global TE derepression coincident with meiotic entry; Antisense TE transcripts rose by 8% over the same interval, which was not significant (p > 0.05). **F.** Scatter plot of log_2_(counts + 1) at E10.5 (x-axis) versus E16.5 (y-axis) for TE families and protein-coding genes, colored by class (genes, black; LINE, blue; SINE, purple; LTR/ERV, red; DNA transposons, orange), with PIWIL1 highlighted in green. TE classes, particularly LTR/ERV and LINE elements, are shifted above the diagonal, indicating preferential TE upregulation at the meiotic stage (two-sided Wilcoxon rank-sum test, adjusted p < 0.05).

We next asked whether induction of this pathway coincided with increased TE transcription. Quantifying TE-derived transcripts with TEtranscripts, we found a significant expansion of sense TE expression at meiotic entry (Fig. 1E; two-tailed Student’s t-test, p < 0.05), accompanied by a smaller, non-significant (8%) increase in antisense TE transcripts from E10.5 to E16.5 (Fig. 1E, p > 0.05). The increase was dominated by the retrotransposon classes LTR/ERV and LINE (Fig. 1F; two-sided Wilcoxon rank-sum test, adjusted p < 0.05). 84.4% of expressed TE families showed higher expression at E16.5, spanning retroelements (CR1, ERVL, ERV1) and DNA transposons (Helitron, hAT) (Table S2). Because ERVs and LINEs are the principal substrates of piRNA-mediated silencing, we focused on these families. The most strongly induced elements were the endogenous retrovirus GGERV11 and the CR1/LINE subfamilies CR1-16_Crp, X7A-LINE, and X7B-LINE (Fig. 2A, DESeq2 Wald test, Benjamini-Hochberg-adjusted, p < 0.05), nominating these as candidate priority targets for silencing. Because whole-ovary measurements can be confounded by the changing germ cell fraction between stages, we sought confirmation at single-cell resolution.

**Figure 2.**
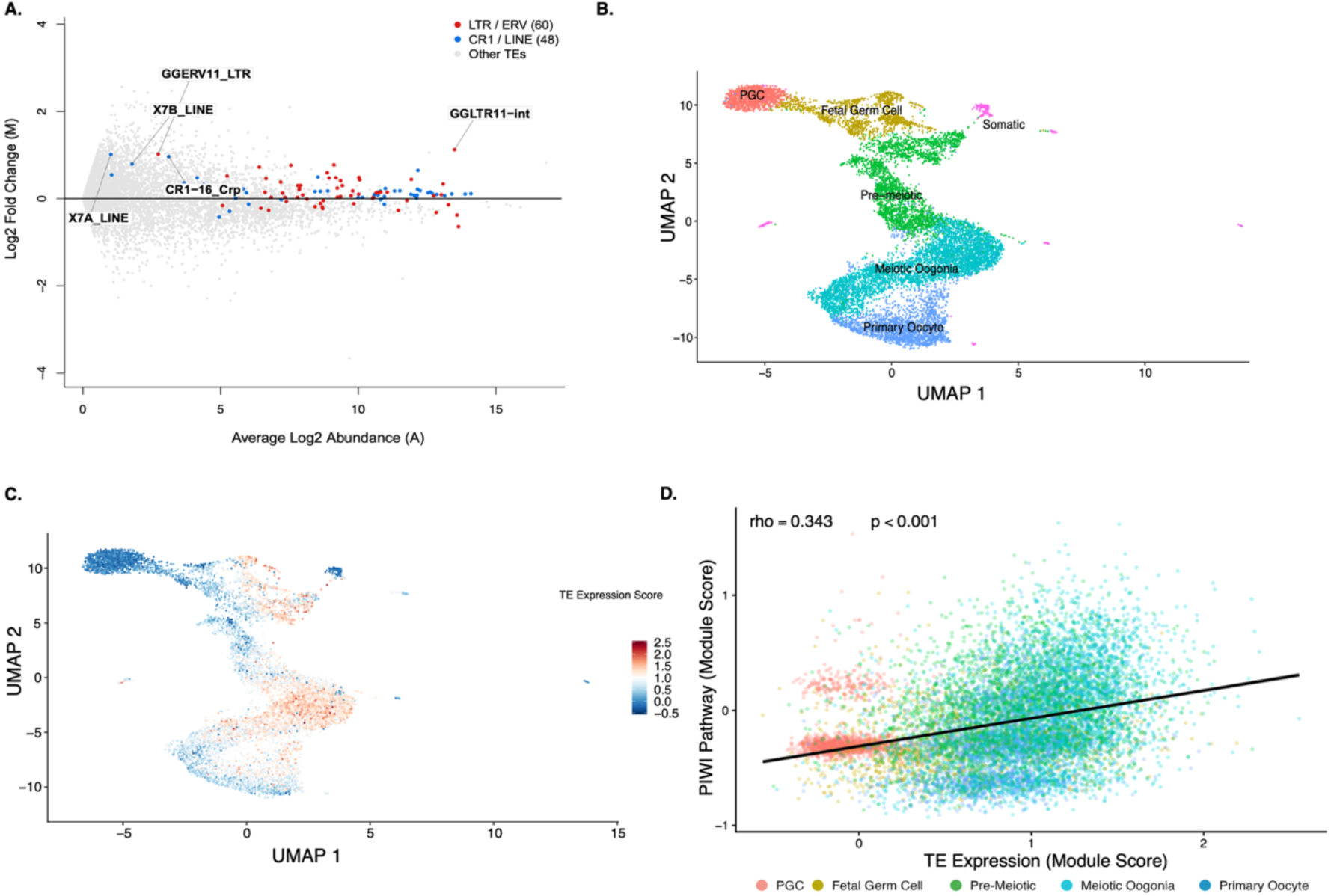
Transposable element derepression tracks with piRNA-pathway activation across single germ cells. **A.** MA plot of TE-family differential expression between E16.5 and E10.5 ovaries (TEtranscripts/DESeq2): log_2_ fold change (M) plotted against average log_2_ abundance (A). LTR/ERV families are shown in red (n = 60 families tested) and CR1/LINE families in blue (n = 48); other TEs are shown in grey. Labeled families are the most strongly induced: GGERV11 (GGLTR11/GGERV11-int), X7B-LINE, X7A-LINE, and CR1-16_Crp. **B.** UMAP of ovarian cells from the published scRNA-seq dataset (PRJNA761874), colored by annotated cell type: PGCs (red), Fetal Germ Cells (olive), Pre-meiotic (green), Meiotic Oogonia (cyan), Primary Oocytes (teal), and residual somatic cells (pink). DAZL⁺ germ cells were subset from this dataset for the analyses in C, D, and Fig. S1A. **C.** UMAP of DAZL⁺ germ cells colored by the global TE expression score. TE expression score increases progressively from PGCs through meiotic stages, peaking in Meiotic Oogonia. **D.** Scatter plot of the global TE expression score (x-axis) versus the PIWI pathway score (y-axis; score of PIWIL1, TDRD1, TDRD5, TDRD9, TDRD12, MYBL1, MOV10L1) across individual germ cells, colored by developmental stage as in B. A moderate positive association is observed (Spearman ρ = 0.343, p < 0.001), consistent with meiosis-coupled co-induction of TE transcription and piRNA-pathway gene expression score.

To resolve whether TE reactivation and pathway induction track together at the cellular level, independently of bulk tissue composition, we analyzed published scRNA-seq of DAZL⁺ germ cells spanning PGCs to primary oocytes (PRJNA761874 (23); Fig. 2B). A global TE-expression score increased progressively along the trajectory, peaked in meiotic oogonia, and declined in primary oocytes (Fig. 2C, Fig. S1A; ANOVA, p < 0.05). Across single germ cells, TE expression score was moderately associated with piRNA-pathway gene expression score, scored from *PIWIL1*, *TDRD1*, *TDRD5*, *TDRD9*, *TDRD12*, *MYBL1*, and *MOV10L1* (*TDRKH*, *M1AP*, and *MAEL* were excluded owing to high single-cell dropout) (Spearman ρ = 0.343, p < 0.001; Fig. 2D). Together, these data show that meiotic entry in the female chicken germline is marked by a coordinated developmental switch, exit from pluripotency, induction of meiotic factors, and PIWIL1-led piRNA-pathway activation, occurring alongside a broad transposon burst led by LTR/ERV and LINE elements. This concurrence of a rising TE threat and an induced piRNA pathway at meiotic entry prompted us to ask whether the cytoplasmic arm of the pathway, ping-pong amplification, is actively engaged at this stage, which we address next.

### The ping-pong cycle is engaged against upregulated transposable elements at meiotic entry

To characterize the small-RNA dynamics of the mitotic-to-meiotic transition, we compared small RNA-seq profiles from E10.5, E16.5, and mature ovaries (n = 3). We observed an increase in the piRNA fraction at E16.5 compared with E10.5, followed by a significant downregulation in mature ovaries (Fig. S1B, Fig. 3A; p < 0.05). TE-derived piRNA sequences upregulated at E16.5 shift the scatter plot distribution above the diagonal, with only a minor downregulated population, and antisense piRNAs of TEs increased sharply, a composition consistent with active piRNA biogenesis (Fig. 3B, two-tailed Student’s t-test, p < 0.05).

**Figure 3.**
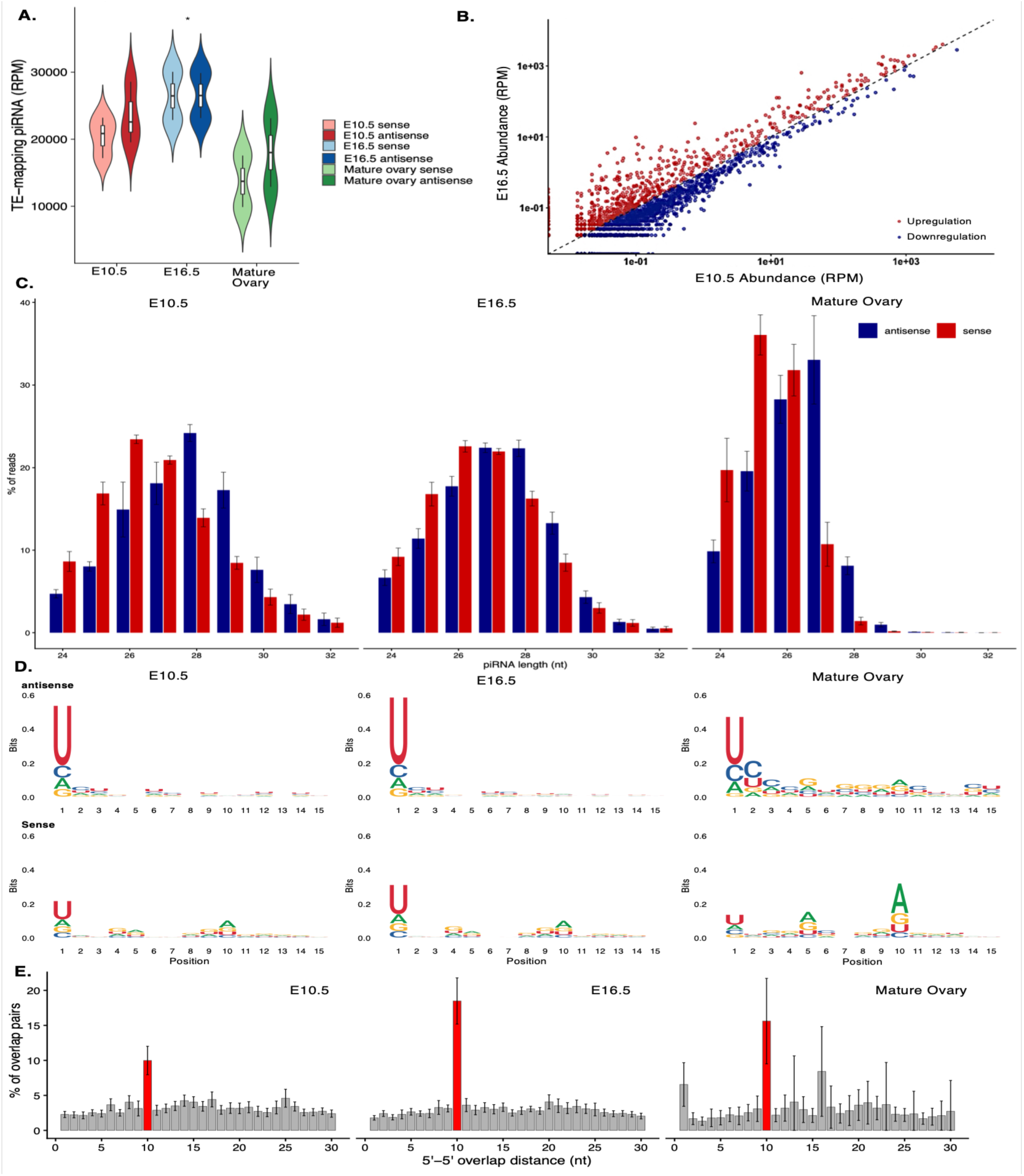
Stage-dependent intensification of piRNA biogenesis and ping-pong amplification at meiotic entry in the chicken ovary. **A.** TE-mapping piRNA abundance (RPM) at E10.5 (mitotic), E16.5 (meiotic entry), and mature ovary, split by strand (light, sense; dark, antisense). Violins show the distribution across replicates; boxplots show median and interquartile range (pairwise two-sided Welch’s t-test, *p < 0.05). **B.** Scatter plot of TE-derived piRNA sequence abundance (RPM, log scale) at E10.5 (x-axis) versus E16.5 (y-axis). Red, sequences more abundant at E16.5 (upregulated); blue, more abundant at E10.5 (downregulated). The predominance of red points above the diagonal reflects higher normalized abundance of many TE-derived piRNA sequences at meiotic entry. **C.** Length distributions (24–32 nt) of sense (red) and antisense (blue) TE-mapping piRNAs at E10.5, E16.5, and mature ovary (% of reads; mean ± SD, n = 3). A canonical 26–30 nt peak is present at all three stages. **D.** Nucleotide composition (positions 1–15; sequence logos, bits) of antisense (top row) and sense (bottom row) TE-mapping piRNAs at each stage. A strong 5ʹ-uridine (1U) bias, the signature of primary piRNA biogenesis, is detected at E16.5 in both strand populations, is markedly weaker at E10.5, and is the least in the mature ovary. The sense population at E16.5 additionally shows a significant position-10 adenine (10A) enrichment, the signature of secondary ping-pong piRNAs (two-sided Welch’s t-test, p < 0.05). **E.** 5ʹ–5ʹ overlap-distance analysis of complementary piRNA pairs mapping to TE loci at each stage; the 10-nt overlap diagnostic of ping-pong amplification is shown in red. A discrete 10-nt peak is present at all three stages, is higher at E16.5 (z = 20.8 ± 6.3) than at E10.5 (z = 7.3 ± 2.5, mean ± SD, n = 3; two-sided Welch’s t-test, p = 0.05, significance threshold z > 3.3), and is lowest in the mature ovary (z = 5.9±4.8, mean ± SD, n = 3; two-sided Welch’s t-test, p < 0.05; significance threshold z > 3.3).

At both E10.5 and E16.5, reads in the mature-piRNA window (26–30 nt; Fig. 3C) carried a 5ʹ-uridine (1U) bias in the antisense and sense mapping populations, the signature of primary piRNAs; the bias was stronger at E16.5 (Fig. 3D; two-sided Welch’s t-test, p < 0.05). Its ping-pong partner, the sense-mapping population, showed the 10-adenine (10A) signature of secondary piRNAs, again stronger at E16.5 compared with E10.5 (Fig. 3D; p < 0.05). Mature-ovary small RNA (n = 3; PRJNA412674) showed the weakest 1U biases of the three stages (Fig. 3A, C and D). To determine whether this program included an active ping-pong cycle, we measured the 5ʹ–5ʹ overlap of complementary piRNA pairs mapping to TE loci and assessed its enrichment by z-score (threshold z > 3.3). A 10-nt overlap peak, the signature of ongoing secondary biogenesis, was present at both fetal stages and was higher at meiotic entry (z = 7.3±2.5 at E10.5 versus 20.8 ±6.3 at E16.5; two-sided Welch’s t-test, p = 0.05, Fig. 3E). Ping-pong amplification is therefore already engaged before meiotic entry and intensifies as meiosis begins. In the mature ovary, a smaller but discrete 10-nt 5ʹ–5ʹ overlap peak persisted compared to meiosis onset (z = 5.9±4.8 compared to E16.5, two-sided Welch’s t- test, p < 0.05; Fig. 3E). The fraction of piRNAs with a 10-nt partner followed the same stage-wise pattern (Fig. S1C, p < 0.05). This was in accordance with the retargeting of ping-pong amplification onto DNA, ERVK, and ERVL elements in mature ovaries, while CR1 elements were more actively targeted at meiosis onset (Fig. S2A, ANOVA, p < 0.05).

We then asked if this tissue-level signal is from the germ-cell compartment. In *DAZL*⁺ germ cells (PRJNA761874, Fig. 2B), *PIWIL1*, *MYBL1*, and *TDRD*-family transcripts were low in PGCs (*NANOG*⁺) and fetal germ cells (*PCNA*⁺), increased through the pre-meiotic stages, peaked in meiotic oogonia (SPO11⁺) at E16.5, and declined in primary oocytes, a transient induction peaked to the meiotic window (Fig. S2B; pre-meiotic and meiotic oogonia compared to other stages, p < 0.05). Meiotic markers (*REC114*, *SPO11*) showed the same sequential rise (Fig. S2B; p < 0.05). Next, we tested whether this pattern is gonad-specific in chicken body tissues and PGCs. Analysis of 21 tissues of chicken organs showed that pathway-gene induction is concentrated in the gonads across the body and tissues (Fig. S2C, Wilcoxon rank-sum test, p < 0.05). Consistent with germline specificity, *PIWIL1* and core silencing-complex (*TDRD*s) transcripts were detected in germ cells (*DAZL*⁺) but not somatic cells (*DAZL*^-^) in independent E17 ovary datasets (Fig. S2D, https://home.kaessmannlab.org/resources) (24).

### Meiosis-associated ping-pong dynamics are conserved in zebrafish

To test whether this program is conserved beyond chicken, we performed equivalent single-cell (GSE173718) and small-RNA analyses in zebrafish ovarian germ cells spanning pre-meiotic to mature stages (Fig. 4A) (25). A global TE-expression score, computed at single-cell resolution, was low in the PGC cluster and increased sharply in meiotic oogonia along the same developmental trajectory used for cell-type annotation (Fig. 4B; ANOVA, p < 0.05), reproducing the coupling between TE expression score and meiotic progression identified in chicken. We then tested for ping-pong amplification at each stage. A significant 10-nt 5ʹ–5ʹ overlap peak was present before meiosis I (3 wpf (weeks post-fertilization) gonad, z = 16±1.4, n = 1, SD is subsampling), at meiosis I (6 wpf ovary, z = 34.8 ± 15.9, n = 5), and in the mature ovary (24 wpf, z = 28.2±6.4, n = 5) (Fig. 4C). Primary (1U, antisense-enriched) and secondary (10A, sense-enriched) piRNA signatures, interpreted as in the chicken analysis above, were detected at all three stages, alongside piRNA size distribution consistent with canonical biogenesis (Table S3). We also observed the same pattern of increase in the meiosis peak and retargeting for the piRNA fraction and for the fraction of piRNAs having a 10-nt partner (Fig. S3A and B, two-sided Welch t-test, *p < 0.05). The peak was present before meiosis I, was highest at meiosis I, and was reduced in the mature ovary, the same stage-wise pattern seen in chicken. Per-TE-family analysis showed the same qualitative retargeting identified in chicken; ping-pong pairs were broadly distributed across nearly all annotated TE families in meiosis, whereas by the mature ovary stage, they narrowed to a restricted subset dominated by LTR-class retrotransposons, LINEs, and DNA transposase elements (Fig. 4D, ANOVA, P < 0.05).

**Figure 4.**
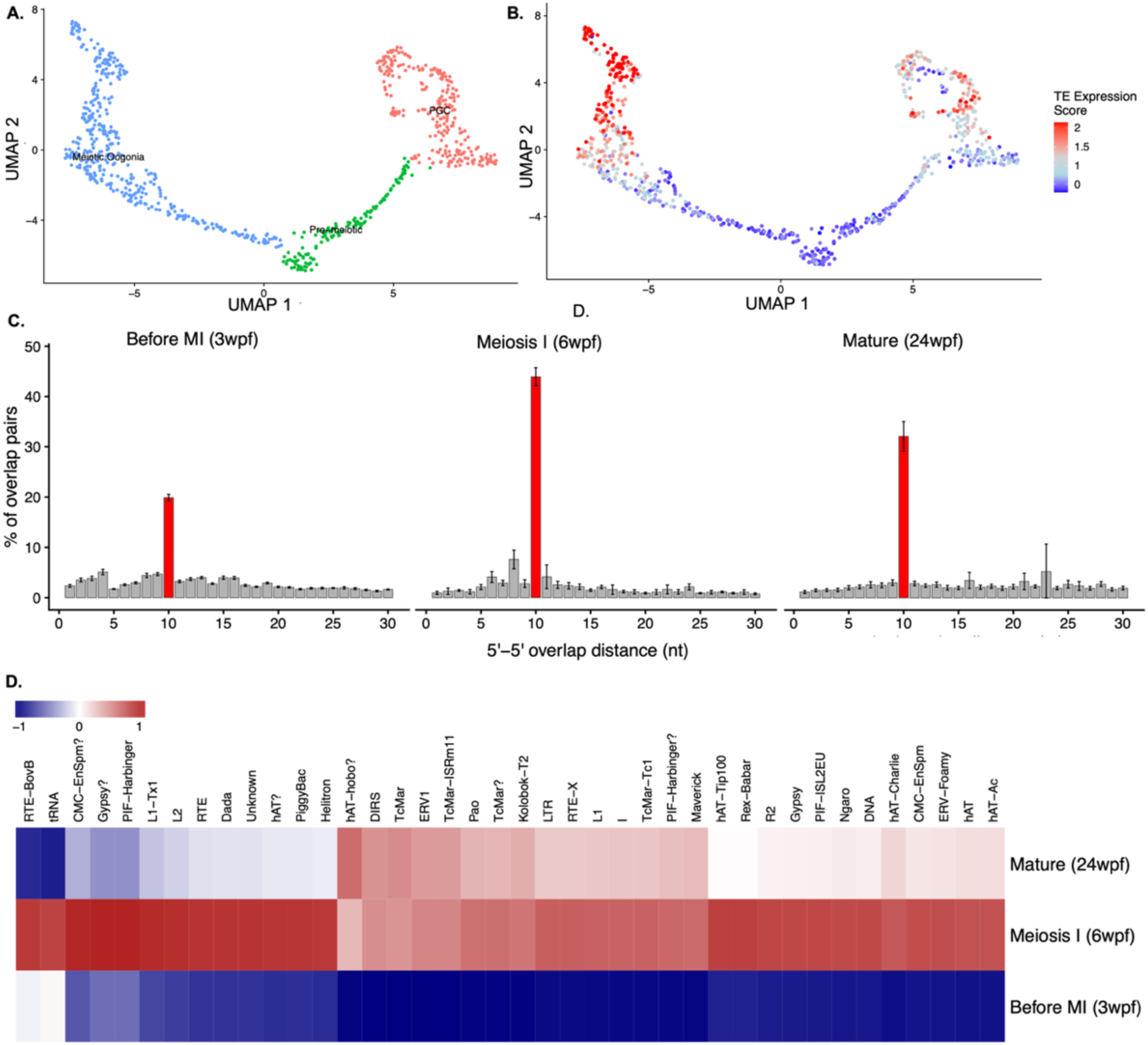
Stage-resolved ping-pong dynamics in the zebrafish germline. **A.** UMAP of zebrafish ovarian germ cells (GSE173718) colored by cell-type annotation: PGC (nanos3), pre-meiotic (rec8), and meiotic oogonia (sycp3). **B.** The same UMAP colored by transposable- element (TE) expression score (blue = low, red = high). The TE score is low in PGCs and rises in the meiotic-oogonia population, indicating that TE expression increases with meiotic progression. **C.** 5ʹ–5ʹ overlap-distance distributions for complementary piRNA pairs mapping to TE loci at three developmental stages (left to right): before meiosis I (3 wpf gonad), meiosis I (6 wpf ovary), and mature ovary (24 wpf). The y-axis shows the percentage of overlap pairs at each distance; the 10-nt overlap (red bar) is the ping-pong signature. Z-scores of the 10-nt peak: z = 16±1.4 (3 wpf; single library, n = 1, SD is subsampling variation), z = 34.8 ± 15.9, (6 wpf; mean ± SD, n = 5), and z = 28.2±6.4 (24 wpf; mean ± SD, n = 5) . The peak is highest at meiosis I and lower before meiosis I and in the mature ovary, the same stage-wise pattern seen in chicken (Fig. 3E).**D.** Heatmap of ping-pong pair abundance per TE family (columns) across the three stages (rows, top to bottom: mature 24 wpf, meiosis I 6 wpf, before meiosis I 3 wpf), scaled per family (blue = low, red = high). Ping-pong pairs are broadly distributed across nearly all annotated TE families at meiosis I, whereas in the mature ovary they are restricted to a subset of families spanning LTR retrotransposons (DIRS, ERV1, Pao, LTR), LINEs (RTE-X, L1, I), and DNA transposons (TcMar-family elements, Kolobok-T2, hAT-Charlie), (ANOVA, P < 0.05).

The two species also differed systematically in absolute z-score magnitude and piRNA population while showing the same pattern at every comparable stage. RepeatMasker annotation showed transposable elements comprise 54.48% of the zebrafish genome (danRer11) versus 9.51% of the chicken genome (galGal6/GRCg6a), a roughly 5.7-fold difference in genomic transposon content (Fig. S3C). This substantially larger TE complement likely accounts for zebrafish’s uniformly higher z-scores and raw pair counts relative to chicken at every developmental stage, since a greater density of genomic TE loci provides more opportunity for ping-pong pairing overall.

### Meiotic induction in cultured PGCs activates primary piRNA biogenesis signatures

To test whether activation of the meiotic program in vitro enhances ping-pong amplification, we treated cultured chicken primordial germ cells (PGCs) with retinoic acid (RA) and BMP2, established inducers of meiotic entry. Treated PGCs upregulated the pre-meiotic marker *STRA8* (Fig. 5A and B; adjusted p < 0.05) (26), confirming induction of the meiotic program, and the piRNA biogenesis factors *PLD6* and *TDRD15* were also induced (Fig. 5B), with weaker upward trends across other pathway components. Global TE expression did not shift relative to an abundance-matched protein-coding background (Fig. 5C; two-sided Wilcoxon rank-sum test, n.s.). ERVK-family elements were nonetheless induced (Fig. 5D; DESeq2 Wald test, Benjamini-Hochberg-adjusted, p < 0.05).

**Figure 5.**
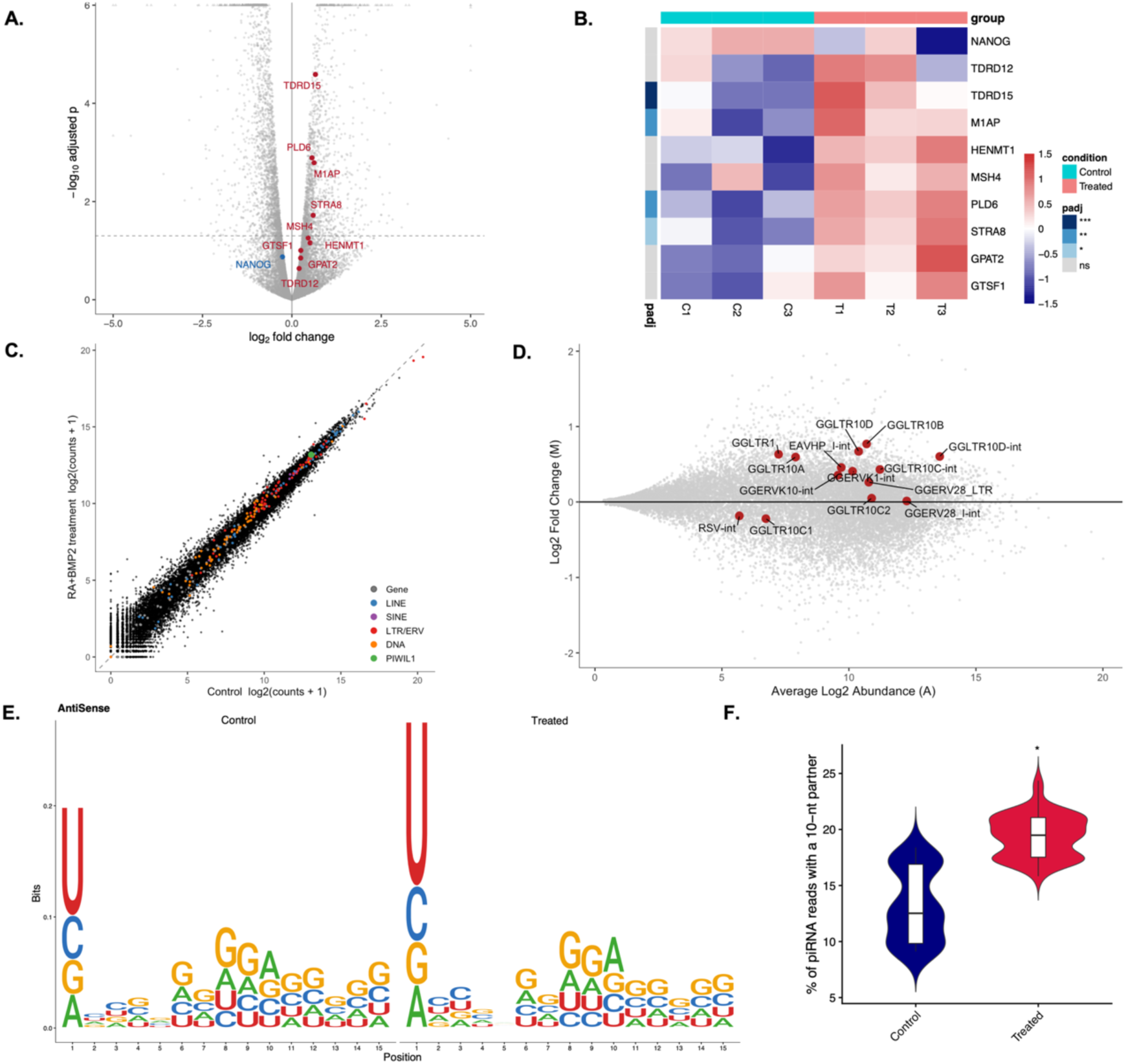
Meiotic induction in cultured chicken PGCs primes piRNA biogenesis and increases ping-pong participation. **A.** Volcano plot of differential gene expression between RA+BMP2-treated and control PGCs (n = 3; DESeq2, Benjamini-Hochberg-adjusted p). Upregulated piRNA-pathway and meiotic genes are labeled in red (TDRD15, PLD6, M1AP, STRA8, MSH4, HENMT1, GPAT2, GTSF1, TDRD12); the pluripotency marker NANOG (blue) is downregulated. **B.** Heatmap of row-scaled normalized expression (z-score) for piRNA-pathway, meiotic, and pluripotency genes in control (C1–C3) and RA+BMP2-treated (T1–T3) PGCs. The left sidebar indicates adjusted p-value significance (***p < 0.001, **p < 0.01, *p < 0.05, ns, not significant). **C.** Scatter plot of log_2_(counts + 1) in treated versus control PGCs for genes (black) and TE families colored by class (LINE, blue; SINE, purple; LTR/ERV, red; DNA, orange (two-sided Wilcoxon rank-sum test, n.s.); PIWIL1 is highlighted in green. Global TE expression did not differ significantly between conditions. **D.** MA plot of TE-family differential expression (TEtranscripts/DESeq2). Significantly differentially expressed families (DESeq2 Wald test, Benjamini-Hochberg-adjusted, p < 0.05) are shown in red and labeled; families are predominantly ERVK-class LTR/ERV elements (GGLTR10, GGERVK, GGERV28, and related internal/LTR subfamilies). **E.** Sequence logos of the first 15 nt of TE-mapping piRNAs in RA+BMP2-treated PGCs compared to non-treated PGC control. The antisense of RA+BMP2-treated PGCs shows a markedly higher amount of 1U bias in antisense strands (two-sided Welch’s t-test, p < 0.05). **F.** Ping-pong participation in control versus treated PGCs, measured as the percentage of piRNA reads with a 10-nt 5ʹ–5ʹ overlap partner (violins with boxplots showing median and interquartile range). Ping-pong participation is higher in treated than in control PGCs (two-sided Welch’s t-test, p < 0.05).

We next profiled small RNAs from the same samples. Although neither the piRNA fraction nor the piRNA size distribution changed detectably (Fig. S4A and B, n.s.), the 5ʹ-uridine (1U) bias of TE-mapping piRNAs strengthened in antisense (Fig. 5E, two-sided Welch’s t-test, p < 0.05), and the fraction of piRNAs with a 10- nt ping-pong partner increased significantly (Fig. 5F; two-sided Welch’s t-test, p < 0.05). The prominence of the 10-nt peak in the 5ʹ–5ʹ overlap-distance distribution, however, did not differ between treated and control PGCs (Fig. S4C; z = 19.2 ± 0.9 versus 19.6 ± 1.9, two-sided Welch’s t-test, n.s.). Induction of the meiotic program in cultured PGCs increased primary piRNA biogenesis signatures and increased ping-pong participation, but it did not measurably sharpen the 10-nt overlap peak. Ping-pong is already active in untreated PGCs, consistent with the pre-meiotic competence seen *in-vivo*, so treatment increased the fraction of reads with a 10-nt partner, while the overlap-enrichment z-score did not change detectably.

## Discussion

In this study, we define a meiosis-coupled genome-defense program in the female chicken germline. Meiotic entry is accompanied by a broad burst of transposable-element (TE) transcription, encompassing 84.4% of TE families and dominated by endogenous retroviruses, together with coordinated induction of the piRNA pathway. PIWIL1 and associated biogenesis factors increase along the trajectory from primordial germ cells (PGCs) to meiotic oogonia. This induction is productive, the piRNA pool expands and is retargeted toward ERV families; the 5ʹ-uridine (1U) bias of antisense piRNAs strengthens alongside a position-10 adenine (10A) signature in their sense partners; and the 10-nt 5ʹ–5ʹ overlap (ping-pong) signature, already detectable before meiotic entry, peaks at meiotic entry. The observation that the same stage-resolved coupling occurs in zebrafish and that its biogenesis arm can be initiated in cultured chicken PGCs by RA and BMP2 suggests that the linkage between meiotic progression and piRNA-mediated defense represents a shared developmental pattern in chicken and zebrafish.

These findings add to growing evidence that the male-centric requirement for piRNAs documented in mice, where loss of the Piwi pathway causes male sterility with little consequence for oogenesis (27, 28), is not representative of vertebrates. In golden hamsters, PIWI proteins and maternally provided piRNAs are essential for the production of functional oocytes (14), and in zebrafish, Piwi/piRNA function is required for germ cell maintenance, differentiation, and meiosis in both sexes (29, 30). The sharp, germ cell-specific induction of the piRNA pathway at female meiotic entry places the chicken expression program closer to the hamster and zebrafish situations than to the mouse. Two comparisons with the male germline sharpen this conclusion. First, the ping-pong z-score at female meiotic entry (z = 20.8 ± 6.3 at E16.5) is comparable to that in the adult testis, though the two were computed with different overlap-scoring schemes and the comparison is qualitative. Second, the developmental pattern differs between the sexes. TE-directed ping-pong in the chicken testis intensifies at pachytene (17), whereas the ovarian signature peaks at meiotic entry, before pachytene, a pre-pachytene pattern closer to that of the mammalian male germline than to that of the chicken male. The heterogametic (ZW) constitution of the female chicken, with its largely unsynapsed sex chromosome pair at pachytene, may impose additional silencing demands, and further comparison with the male (ZZ) germline will be informative in this regard.

The coupling of a TE burst to meiotic entry is best explained by the epigenetic context in which it occurs. Germline development entails genome-wide erasure of DNA methylation, opening a window of vulnerability to TE derepression that is countered by piRNA- and chromatin-based safeguards; meiotic entry adds replication, programmed double-strand breaks, and large-scale chromatin reorganization to this already permissive landscape. The positive correlation between TE expression and PIWI-pathway scores across single germ cells (Spearman ρ = 0.343) is consistent with a defense that scales with threat. Two observations indicate that ERV derepression in particular is an intrinsic feature of the pre-meiotic program rather than a product of the ovarian niche. ERV families dominate the *in-vivo* TE upregulation, and ERVK- class elements were the families selectively induced by RA+BMP2 in cultured PGCs.

A second principle emerging from our stage-resolved analysis is that a ping-pong signature was detectable at all sampled stages, while amplification intensified at meiosis and was stage-specifically targeted. A discrete 10-nt overlap peak was present at every stage in both species, including pre-meiotic stages, indicating that the machinery is assembled and operative before meiosis. What changes at meiotic entry is the intensity of the signature, maximal z-scores at E16.5 in chicken and at 6 wpf in zebrafish, and its target repertoire. Ping-pong pairs are distributed broadly across nearly all annotated TE families at meiosis and subsequently retarget to a restricted subset in the mature ovary, especially in zebrafish (ERVL/ERVK/DNA families in chicken; LTR, LINE, and DNA-class families in zebrafish). This retargeting mirrors the adaptive logic described in the testis, where ping-pong amplification preferentially engages transcriptionally active TE families and is absent from inactive ones, and suggests a transition from broad surveillance during the high-risk meiotic window to focused, family-specific maintenance in the adult ovary.

The in vitro induction experiment delineates which components of this program meiotic signaling alone can engage. RA+BMP2 treatment induced STRA8, upregulated piRNA biogenesis factors (PLD6, TDRD15), strengthened the 1U bias of the antisense-mapping population, selectively induced ERVK families, and significantly increased the fraction of piRNAs engaged in 10-nt pairs. Yet the prominence of the 10-nt peak in the overlap-distance distribution was unchanged (z = 19.2 ± 0.9 versus 19.6 ± 1.9, n.s.). Because these metrics report distinct quantities, the fraction of reads participating in ping-pong pairs versus the sharpness of the 10-nt register, their dissociation indicates that pre-meiotic induction expands the pool of ping-pong- engaged piRNAs without concentrating pairing at the canonical register. Full amplification of the magnitude observed *in-vivo* may require the complete meiotic chromatin environment, organized nuage, the full complement of ping-pong helicases (*MOV10L1*, *TDRD9*), or the far higher TE substrate load of the E16.5 ovary. Cultured PGCs thus provide a tractable system for separating the signaling-driven components of the program from those licensed by the meiotic state itself.

Finally, the cross-species comparison reveals that ping-pong output scales with genomic TE content. The zebrafish genome is 54.5% TE, versus 9.5% in chicken (danRer11 and galGal6/GRCg6a RepeatMasker annotations), and zebrafish exhibited uniformly higher z-scores and pair counts at every comparable stage while following the same qualitative trajectory. This scaling might support a supply-side model in which the density of transcribed TE loci determines the opportunity for ping-pong pairing (31).

## Materials and Methods

### Sample Collection

Fertilized Lohmann *Gallus gallus* eggs were obtained from a commercial breeder and incubated at 37.5 °C with approximately 60% relative humidity under standard conditions. Left ovaries, the functional gonad in birds (32) were dissected from female embryos at embryonic day 10.5 and embryonic day 16.5 under a stereomicroscope and validated based on the Hamburger-Hamilton (HH) stage; sex was determined as previously shown, n = 3 per biological group, independent embryos (33). Tissues were isolated under sterile, RNase-free conditions, immediately flash-frozen in liquid nitrogen, and stored at −80 °C until RNA extraction. All handling and surgical procedures adhered to local ethics committee guidelines, and animal experiments were conducted with approval of Uppsala University.

### RNA extraction

Total RNA was extracted using TRIzol™ Reagent (Thermo Fisher Scientific) according to the manufacturer’s instructions. RNA integrity was assessed using an Agilent Bioanalyzer 2100, and only samples with RNA integrity numbers (RIN) greater than 8 were used for downstream applications. RNA concentration was quantified by Qubit 2.0 fluorometer (Qubit RNA HS Assay Kit, Thermo Fisher Scientific).

### Library Preparation and Sequencing

For small RNA sequencing (n = 3 per group), libraries were prepared using the NEBNext® Multiplex Small RNA Library Prep Set for Illumina® (Set 1; Cat. No. E7300) following the manufacturer’s protocol. Briefly, 3ʹ and 5ʹ adapters were ligated to small RNAs, followed by reverse transcription and PCR amplification. Libraries were constructed, and small RNA fragments ranging from 18–40 nucleotides were size-selected. Libraries were sequenced on an Illumina NovaSeq X Plus platform using single-end 50 bp (SE50) chemistry. For bulk RNA sequencing (n = 3, per group), strand-specific libraries were generated using the NEBNext Ultra II Directional RNA Library Prep Kit for Illumina® following poly(A) mRNA enrichment. Briefly, polyadenylated RNA was isolated using oligo(dT) magnetic beads, fragmented, and reverse-transcribed to generate strand-specific cDNA libraries, followed by end repair, adapter ligation, and PCR enrichment. Libraries were sequenced on an Illumina NovaSeq X Plus platform to produce paired-end 150 bp (PE150) reads. To eliminate potential batch effects, all biological replicates (n = 3 per stage, same sample used for both types of sequencing) for both bulk strand-specific RNA-seq and small RNA-seq were processed in parallel.

### Quantitative Real-Time PCR (qRT–PCR)

Total RNA was reverse-transcribed into complementary DNA (cDNA) using the RevertAid First Strand cDNA Synthesis Kit (Thermo Fisher Scientific) according to the manufacturer’s instructions. Quantitative real-time PCR (qRT–PCR) was performed using PowerUp™ SYBR™ Green Master Mix (Thermo Fisher Scientific) on a QuantStudio 6 Flex (Thermo Fisher Scientific). Each reaction was carried out in a final volume of 20 µL and included 10 ng of cDNA. Thermal cycling conditions consisted of an initial activation step followed by amplification cycles according to the manufacturer’s recommendations (50 °C/2 min → 95 °C/2 min → 40× 95 °C 15 s, 60 °C 60 s). Melt curve analysis was performed at the end of each run to confirm amplification specificity. Gene expression levels were normalized to *GAPDH* and *ACTB* (geometric mean), and relative expression was calculated using the 2⁻ΔΔCt method. The qRT-PCR results were analyzed using a two-tailed Student’s t-test. All reactions were performed with three technical replicates, and each group had three biological replicates; primers are listed in Table S4.

### PGC Cell Culture and Meiosis Conditions

Female chicken primordial germ cells were cultivated and maintained in feeder-free culture in a defined medium, as described previously (33). Briefly, defined medium was prepared from a PGC basal medium composed of DMEM (high glucose, no glutamine, no calcium; Thermo Fisher Scientific, Cat# 21068028) adjusted to a final osmolarity of 250 mOsm with distilled water and supplemented with 2.0 mM GlutaMAX, 1× non-essential amino acids (NEAA), 1× penicillin/streptomycin, 0.1 mM β-mercaptoethanol, 1× nucleosides, 0.4 mM sodium pyruvate, 0.1 mg/mL sodium heparin (Sigma-Aldrich), and 0.15 mM CaCl_2_. This basal medium was further supplemented with 0.2% (w/v) ovalbumin (Sigma-Aldrich), 10 µg/mL ovotransferrin (Sigma-Aldrich), 1× B-27 supplement, 30 ng/mL human Activin A (*E. coli*-derived; PeproTech), and 5 ng/mL bFGF (*E. coli*-derived; PeproTech). Cells were maintained at 37 °C in a humidified atmosphere of 5% CO_2_. For meiotic induction, PGCs cultured in defined medium were treated for 7 days with 100 nM RA plus 300 ng/mL bone morphogenetic protein 2 (BMP2-recombinant; PeproTech). Retinoic acid (RA, Thermo Fisher Scientific) was prepared in DMSO; control cultures received an equivalent volume of vehicle. The medium and treatments were refreshed every 2 days, and cells were harvested on day 7 for directional mRNA-seq and small RNA-seq.

### Bioinformatics Analysis

#### Bulk RNA-seq Processing and Differential Expression

Raw sequencing reads were quality-checked using FastQC (v0.11.9, RRID:SCR_014583) and MultiQC (v1.14, RRID:SCR_014982). Adapters and low-quality bases were trimmed using Trim Galore! (v0.6.7, RRID:SCR_011847). Trimmed reads were aligned to the chicken reference genome (galGal6/GRCg6a) using STAR (v2.7.10, RRID:SCR_004463) (34) with strand-specific parameters. Gene-level quantification was performed with featureCounts from the Subread package (v2.0.3, RRID:SCR_012919), using the GRCg6a GTF annotation. Differential gene expression was analysed with DESeq2 (v1.50.2, RRID:SCR_000154) in R, with developmental stage as the design variable. For the PGC meiotic-induction experiment, differential expression was tested with treatment as the design variable. Counts were normalized using the median-of-ratios method in DESeq2, and a variance-stabilizing transformation (VST) was applied for visualization.

#### Transposable Element (TE) Quantification

To quantify the expression of transposable elements, we utilized the TEtranscripts (v2.2.3) (35) pipeline. Multi-mapping reads were reassigned to TE families using the expectation-maximization algorithm implemented in TEtranscripts. We used the RepeatMasker annotation for GRCg6a (RepeatMasker, RRID:SCR_012954, GRCg6a, GCA_000002315.5, combined Dfam_Consensus (release 20170127) and RepBase (release 20170127) repeat libraries) to define TE loci (http://www.repeatmasker.org). TE-family counts from TEtranscripts were tested for differential expression using DESeq2 to identify TE families significantly upregulated at the E16.5 meiotic transition.

#### Functional Enrichment Analysis

To characterize the biological pathways associated with the E10.5 to E16.5 transition, Gene Ontology (GO) enrichment analysis was performed on significantly upregulated genes using the clusterProfiler (v4.6, RRID:SCR_016884) package in R, with all expressed genes as the background universe, using the org.Gg.eg.db annotation; Biological Process terms were tested (36). p-values were adjusted for multiple testing using the Benjamini-Hochberg method.

#### Small RNA-seq Preprocessing and piRNA Identification

For small RNA analysis of all three groups of chicken, adapters were removed using Cutadapt (v5.2, RRID: SCR_011841) (37), and reads were filtered by length to retain sequences between 18–40 nucleotides. After removing reads mapping to structural RNAs (rRNA, tRNA, snRNA, and snoRNA) using Bowtie (v1.3.1)(38), the remaining reads were aligned to the chicken genome (GRCg6a) with STAR (v2.7.10). piRNAs were defined as genome-matching reads of 26–30 nt; length distributions are plotted over 24–32 nt to show the peak and its boundaries. Multi-mapping reads were retained, with all reported alignments kept, to avoid discarding repeat-derived piRNAs, which constitute the majority of the piRNA pool. The 5ʹ nucleotide frequency (1U/10A bias) was visualized using WebLogo (v3.7, RRID:SCR_010236) (39).

#### Small RNA classification

Raw reads were adapter-trimmed with cutadapt (v5.2), retaining inserts of 18–40 nt with a quality cutoff of Q20 and discarding reads in which no adapter was detected. Trimmed reads were collapsed to unique sequences, with read counts retained in the FASTA header. Small RNAs were assigned to classes by sequential alignment with Bowtie (v 1.3.1, RRID:SCR_005476) against a hierarchy of reference sets, each read being assigned to the first class it matched and removed from subsequent rounds. The order was: miRNA, miRBase (release 23) hairpin sequences, sense-strand only, ≤1 mismatch; rsRNA, rRNA and Mt_rRNA transcripts from Ensembl GRCg6a ncRNA (RRID:SCR_002344, release 105), ≤2 mismatches; tsRNA, GtRNAdb tRNA genes with a 3ʹ CCA appended, ≤2 mismatches; sncRNA, snoRNA, snRNA, scaRNA, misc_RNA, Mt_tRNA, sRNA and ribozyme transcripts from the same Ensembl ncRNA set, sense-strand only, ≤2 mismatches; remaining reads were aligned to the galGal6 genome (≤1 mismatch), and genome-matching reads of 26–30 nt were classified as piRNA; mRNA, reads outside the piRNA length window, together with genome-unmapped reads, aligned to Ensembl GRCg6a cDNA, sense-strand only, ≤2 mismatches; lncRNA, Ensembl lncRNA transcripts, sense-strand only, ≤2 mismatches; all remaining reads were designated other.

For each library, reads in each class were tallied per read length and expressed as a percentage of total classified reads. Length distributions are shown. Classification was for composition only.

#### piRNAs, Ping-Pong Signature, and z-Score Calculation

Ping-pong signatures were detected by analyzing 5ʹ-to-5ʹ overlap distances between sense and antisense piRNAs mapping to the same TE locus. All reported alignments were retained at full weight, since repeat- derived piRNAs constitute the majority of the piRNA pool and discarding multi-mappers would preferentially remove reads from the high-copy families of interest. TE-mapping piRNA reads were subsampled to 100,000 per library, repeated 25 times and averaged, to make z-scores comparable across sequencing depths. The frequency of each overlap distance was calculated, and the ping-pong z-score was computed as: Z = (n_10_ − μ)/σ
where n_10_ is the observed frequency of 10-nt overlaps, and μ and σ are the mean and standard deviation of overlap frequencies across positions 1–9 and 11–20 nt (excluding position 10) (40). A z-score > 3.3 was considered evidence of significant ping-pong amplification (17). Reported values are mean ± SD across biological replicates, except for the 3 wpf time point of the zebrafish.

#### scRNA-seq Analysis

Published single-cell RNA-seq data spanning chicken ovarian development from E2.5 to one week post-hatch were obtained from the NCBI SRA (PRJNA761874). Mapping was done with STAR (STARsolo, v2.7.10) (GRCg6a), and raw count matrices were processed using Seurat (v4.3). Clusters were annotated based on the expression of canonical marker genes visualized as feature plots on the UMAP embedding. PGCs were identified by high expression of *NANOG*; Fetal Germ Cells by DAZL positivity combined with expression of the proliferation markers *PCNA* and *MCM2*; Pre-meiotic cells by the onset of *STRA8* and *REC114* expression marking meiotic commitment; Meiotic Oogonia by strong co-expression of *SPO11* and *SYCP2L* reflecting active meiotic double-strand break formation and synapsis; and Primary Oocytes by expression of the post-meiotic oocyte identity markers *LHX8* and *GDF9*. Cells that did not match any germ-cell marker were designated somatic/other and excluded from downstream analysis. Expression dynamics of piRNA pathway genes (*PIWIL1*, *MYBL1*, *TDRD1*, *TDRD5*, *TDRD9*, *TDRD12 and MOV10L1*), meiotic markers (*STRA8*, *SPO11*), and pluripotency factors (*NANOG*, *LIN28A*) were visualized across cell type stages using dot plots and feature plots. Differential expression was assessed using the Wilcoxon rank-sum test implemented in Seurat’s FindMarkers function, with Bonferroni-corrected p-values < 0.05 considered significant. TE loci were defined using the RepeatMasker annotation for GRCg6a (GCA_000002315.5; combined Dfam_Consensus and RepBase libraries, release 20170127). Per-cell TE counts were obtained with scTE (41), a TE expression score (all TE features) and a PIWI-pathway score (PIWIL1, TDRD1, TDRD5, TDRD9, TDRD12, MYBL1, MOV10L1) were calculated with Seurat AddModuleScore, and their association was tested by Spearman correlation using cor.test in R.

#### Zebrafish comparative analysis

To test conservation, we analyzed published zebrafish (*Danio rerio*) single-cell (GSE173718, (42)) and small RNA-seq (PRJNA1483234) data deposited in SRA using the same pipelines (GRCz11; STARsolo/Seurat for single-cell, and the chicken small-RNA/ping-pong workflow). Germ-cell clusters were annotated with established zebrafish markers (*nanos3* (PGC)/ *foxl2l*+*rec8* (Pre-meiotic oogonia)/ *sycp3* (Meiotic oogonia)) (42). Ping-pong z-scores were computed per library and are reported as mean ± SD across n = 5 biological replicates for the 6 wpf (Meiosis I) and 24 wpf (mature) stages; the 3 wpf pre-meiotic stage was represented by a single sample (n = 1).

#### RNA-seq expression analysis

To characterize the tissue-specific expression profiles of the candidate genes (Fig. S2C), we used the publicly available bulk RNA-sequencing dataset generated by the ChickenGTEx project (PMID: 40200121). The ChickenGTEx resource integrates transcriptomic data from thousands of RNA-seq samples representing a broad range of tissues and chicken breeds. Breed information for the ChickenGTEx samples was further matched to the large whole-genome sequencing (WGS) dataset used in our previous study (PMID: 42268884). Samples from wild Red junglefowls were not included in our further analysis.

Processed gene-expression data, quantified as transcripts per million (TPM), were obtained from the ChickenGTEx dataset. For each candidate gene, TPM values were extracted across the available tissues to characterize its tissue-specific expression pattern. Expression values were retained as reported in the original ChickenGTEx dataset without additional normalization. The resulting TPM values were summarized and visualized across tissues to compare the expression profiles of the candidate genes.

## Statistical Analysis

All analyses were performed in R (v4.3.0); n denotes independent biological replicates, data are mean ± SD, and two-sided p < 0.05 was considered significant. Differential expression of genes and TE families was assessed with DESeq2 (Wald test, Benjamini-Hochberg-adjusted p; genes at adjusted p < 0.05 and |log_2_FC| ≥ 1). Class-level TE shifts were tested by two-sided Wilcoxon rank-sum test on per-feature residuals from the diagonal against an abundance-matched protein-coding background, with Benjamini-Hochberg correction across classes. All per-library small RNA metrics, piRNA fraction, length distribution, 1U and 10A nucleotide bias, ping-pong participation, and 5ʹ–5ʹ overlap z-score were computed per biological replicate and compared by two-sided Welch’s t-test (pairwise for the three chicken stages); bootstrap resampling was used only for intervals on individual replicate estimates, not for testing. Ping-pong within a library was called significant at z > 3.3. qRT–PCR used unpaired two-tailed Student’s t-tests. Gene Ontology p-values (clusterProfiler) were Benjamini-Hochberg-adjusted. Single-cell differential expression used the Wilcoxon rank-sum test with Bonferroni correction (Seurat); per-stage TE-score differences were assessed by ANOVA, and TE load versus PIWI-pathway score by Spearman’s rank correlation.

## Data Availability

The raw sequencing data generated in this study (bulk strand-specific RNA-seq and small RNA-seq of chicken embryonic ovaries) have been deposited in the European Nucleotide Archive (ENA) under project accession PRJEB110625. Publicly available datasets re-analyzed in this study include chicken ovarian single-cell RNA-seq (PRJNA761874), zebrafish single-cell RNA-seq (GSE173718), and chicken mature-ovary small RNA-seq (PRJNA412674). Zebrafish small RNA-seq data were provided by, and used with permission from, Igor Szczepan Babiak (Nord University, Norway, PRJNA1483234). Processed sequence count matrices and ping-pong-related tables are provided in the Supplementary Data.

## Acknowledgments

This work was funded by grants awarded to S.Y.T. from the Swedish Research Council for Sustainable Development (Formas; 2023-01396) and the Carl Trygger Foundation (CTS 24:3578), and to A.F. from the Swedish Research Council (VR; 2025-05129), the Swedish Research Council for Sustainable Development (Formas; 2021-00513), and the Carl Trygger Foundation (CTS 24:3571). Claude Opus 4.8 (Anthropic) and ChatGPT GPT-5.5 (OpenAI) were used to assist with language editing and refinement of the manuscript, as well as optimisation and debugging of custom analysis code. The authors retain full responsibility for all scientific content, interpretation of the data, and conclusions.

## Author Contributions

M.H.G. and A.F. conceived and designed the study. M.H.G., A.F., and L.T. developed the interpretation of the results. A.F. supervised the project. M.H.G., S.Y.T., A.N., C.M. and S.C. performed the experiments.

M.H.G. and C.M. performed the data curation, formal analysis, and visualization with input from A.F. and L.T. M.H.G. drafted the manuscript, with contributions from all authors.

## Competing Interest Statement

The authors declare no competing interests.

## Supplementary Figures and Tables

**Figure S1.**
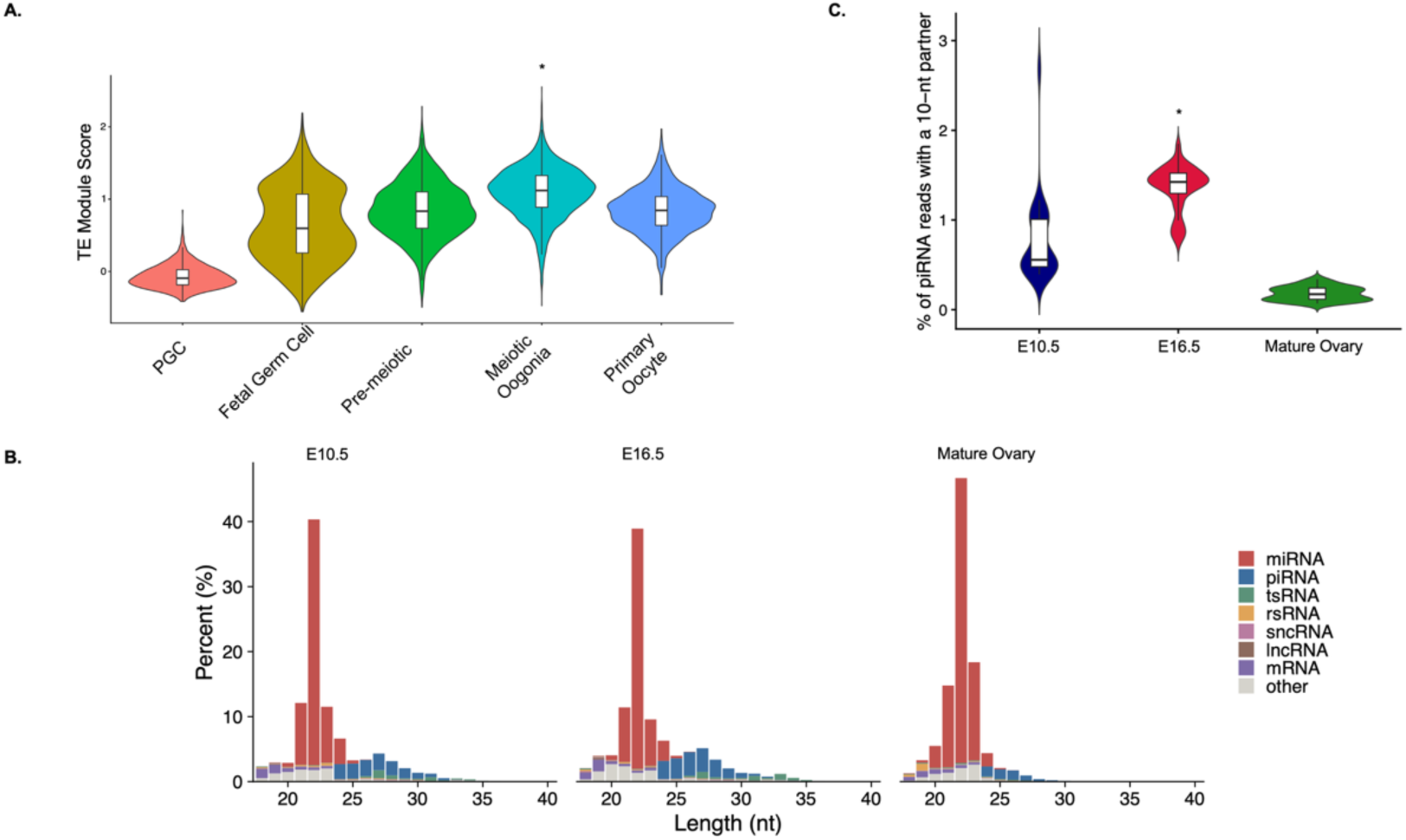
Transposable element expression rises with meiotic progression and coincides with activation of the piRNA ping-pong pathway in the chicken ovaries and primordial germ cells. **A.** Violin plots of per-cell transposable element (TE) expression scores across annotated germ cell populations of the embryonic ovary (PGC, primordial germ cells; Fetal Germ Cell; Pre-meiotic; Meiotic Oogonia; Primary Oocyte). TE expression scores increase with meiotic progression, peaking in meiotic oogonia. Boxes indicate the interquartile range; horizontal lines indicate medians. ANOVA, *P < 0.05. **B.** Length distributions of small RNA-seq reads from whole ovaries at E10.5, E16.5, and mature ovary, colored by RNA biotype. The piRNA-sized fraction (24–32 nt, blue) is markedly expanded at E16.5 relative to E10.5, consistent with developmental induction of piRNA biogenesis around meiotic entry (two-sided Welch’s t-test, *p < 0.05). **C.** Ping-pong amplification signature, quantified as the percentage of piRNA reads with a 10-nt 5ʹ-overlap partner, at E10.5, E16.5, and mature ovary. The signature peaks at E16.5, coincident with peak TE expression, and declines in the mature ovary. Boxes indicate the interquartile range; horizontal lines indicate medians. *P < 0.05, two-sided Welch’s t-test.

**Figure S2.**
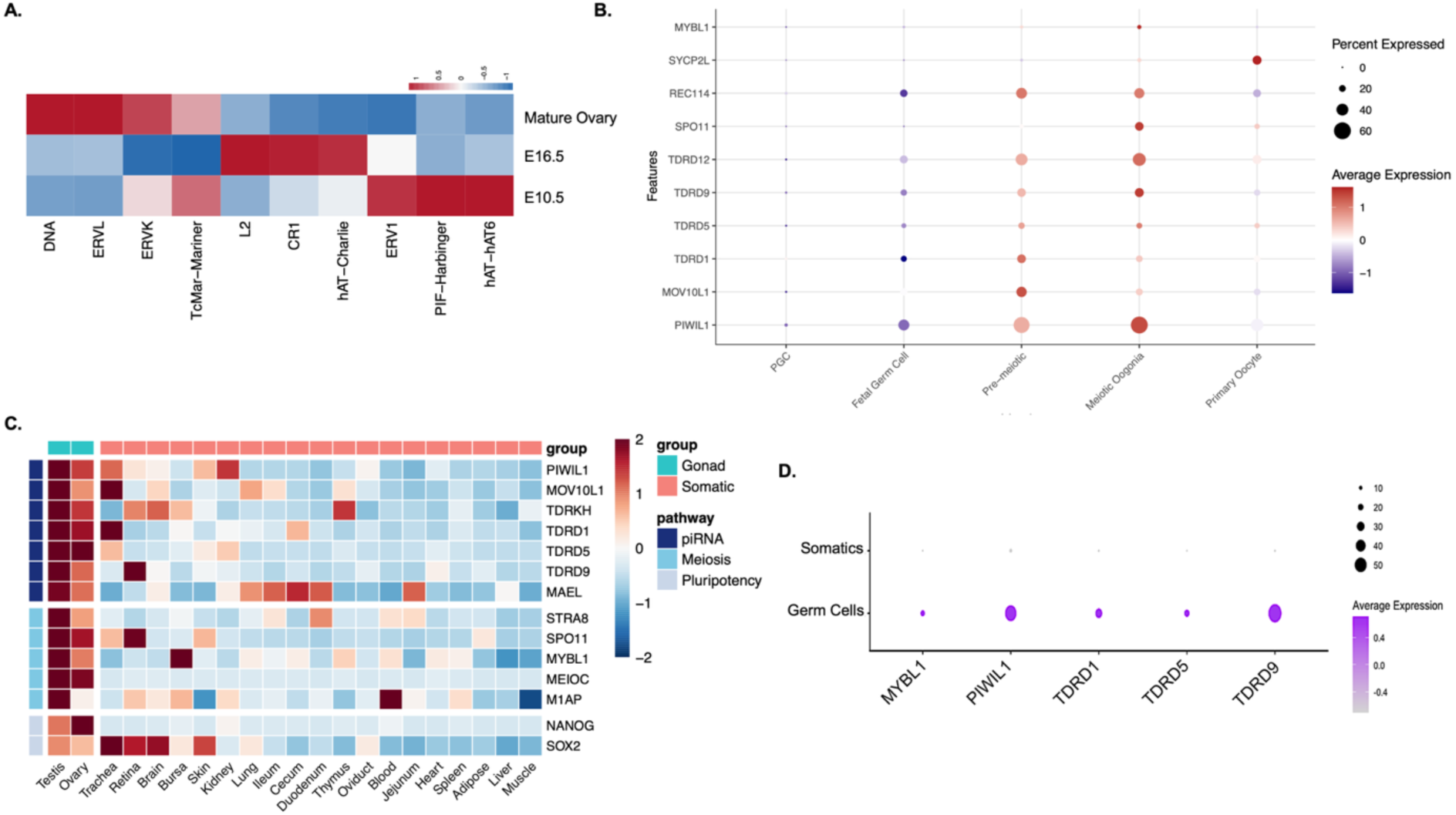
Germ cell-specific piRNA pathway activation and TE repertoire dynamics during the mitotic-to- meiotic transition in the chicken ovary. **A.** Heatmap of per-family ping-pong signal (row z-score; blue = low, red = high) across TE families in whole ovary at E10.5, E16.5, and mature ovary. Families targeted by ping-pong shift between stages, with CR1 elements most active at meiotic entry and DNA and ERVL elements predominating in the mature ovary (ANOVA, P < 0.05). **B.** Dot plot of meiotic marker (MYBL1, SYCP2L, REC114, SPO11) and piRNA pathway (PIWIL1, MOV10L1, TDRD1, TDRD5, TDRD9, TDRD12) genes (rows) across germ cell stages (columns). Color intensity indicates scaled average expression (blue = low, red = high); dot size indicates the percentage of expressing cells. piRNA pathway genes are induced at the pre-meiotic stage and peak in meiotic oogonia (adjusted P < 0.05, Bonferroni-corrected two-sided Wilcoxon rank-sum test). **C.** Heatmap of row-scaled normalized expression (row z-score; blue = low, red = high) for piRNA-pathway, meiotic and pluripotency genes across 21 chicken tissues. Rows are grouped by pathway (left annotation); columns by gonadal versus somatic tissue (top annotation). piRNA-pathway genes are highest in testis and ovary. Wilcoxon rank-sum test, p < 0.05. **D.** Dot plot of piRNA pathway and meiotic marker genes in somatic versus germ cells at E17 (DAZL^+^ versus DAZL^-^). Color intensity indicates average expression (purple gradient); dot size indicates the percentage of expressing cells. All genes shown are restricted to, or markedly enriched in, germ cells, confirming the germline specificity of piRNA pathway activation.

**Figure S3.**
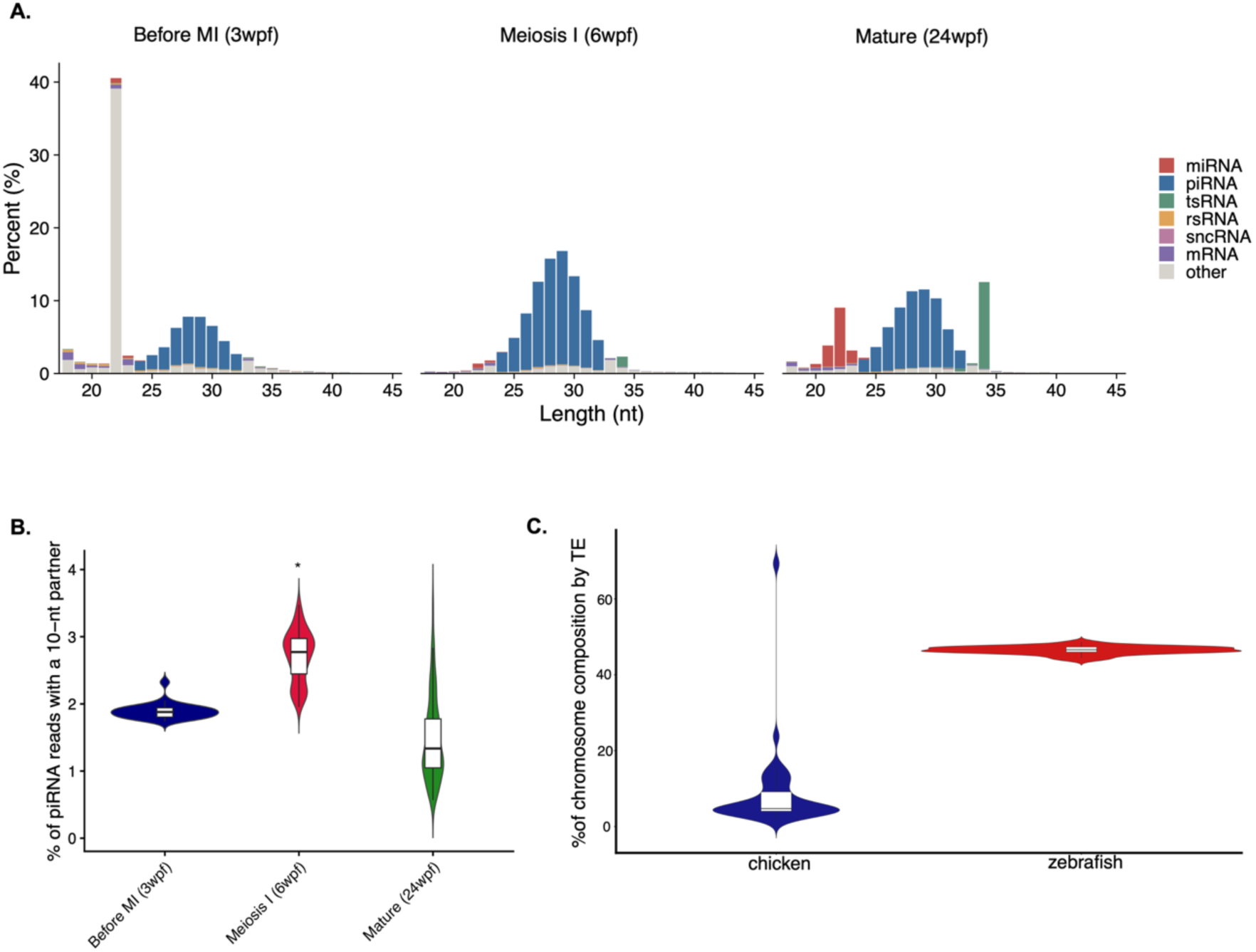
piRNA pool composition and ping-pong participation across zebrafish oogenesis, and genomic TE content in the two species. **A.** Small RNA length distributions (18–40 nt; % of reads) in zebrafish gonad before meiosis I (3 wpf), at meiosis I (6 wpf), and in the mature ovary (24 wpf), stacked by RNA biotype (miRNA, piRNA, tsRNA, rsRNA, sncRNA, mRNA, other). The piRNA-sized fraction (blue, 24–32 nt) is a minor component before meiosis I, where reads are dominated by a 22-nt non-piRNA population, and becomes the predominant biotype at meiosis I and in the mature ovary. **B.** Ping-pong participation across the same three stages, measured as the percentage of piRNA reads with a 10-nt 5ʹ–5ʹ overlap partner. Boxplots show median and interquartile range. Participation is highest at meiosis I and lower before meiosis I and in the mature ovary (two-sided Welch t-test, *p < 0.05). **C.** Percentage of each chromosome annotated as transposable element in chicken (galGal6/GRCg6a) and zebrafish (danRer11). Violins show the distribution across chromosomes; boxplots show median and interquartile range. TEs occupy a substantially larger fraction of the zebrafish genome (54.48% genome-wide) than of the chicken genome (9.51%), a roughly 5.7-fold difference in genomic transposon content.

**Figure S4.**
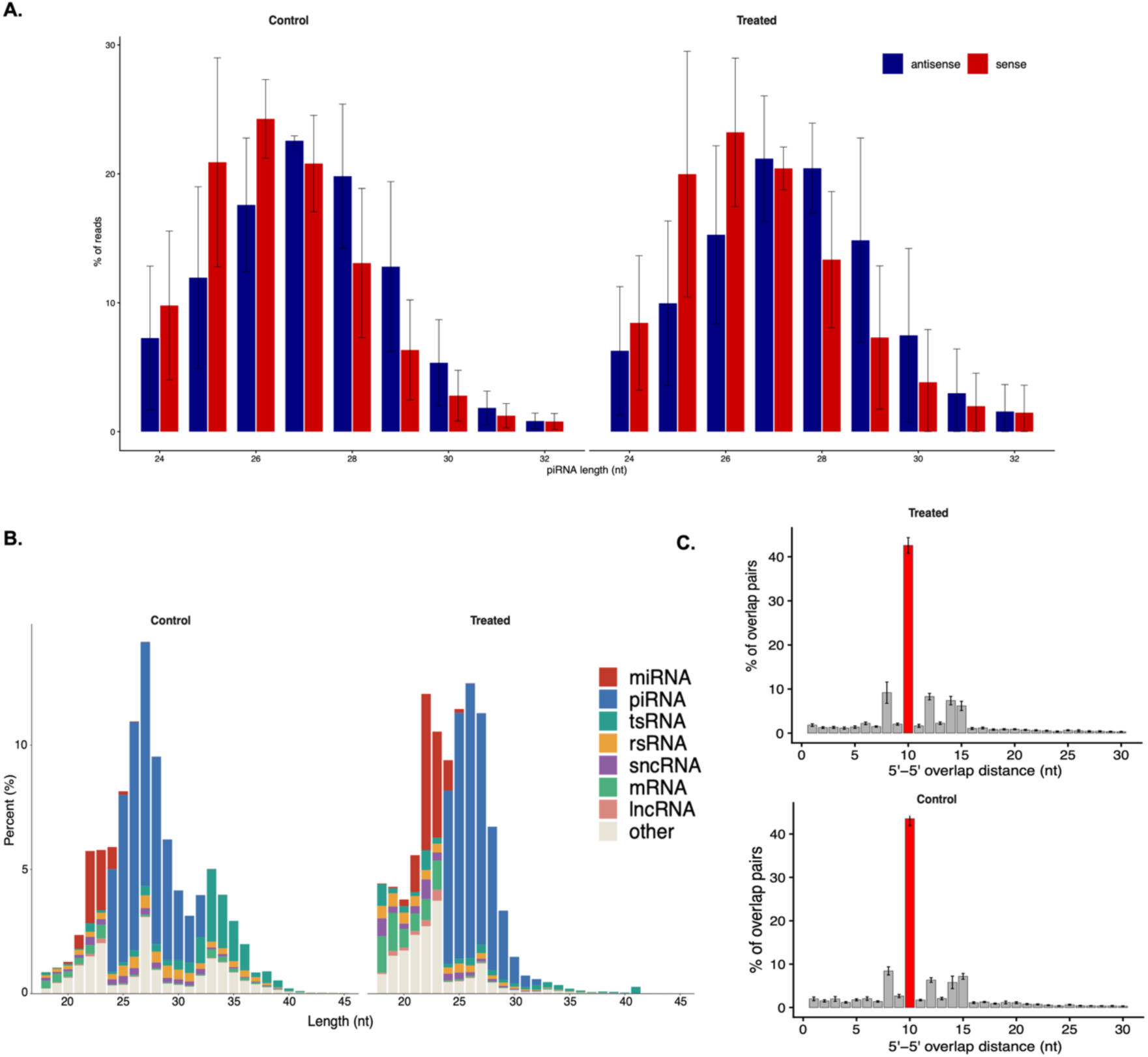
piRNA pool composition and 10-nt overlap strength during meiotic induction of cultured chicken PGCs. A. Length distribution of piRNA reads (26–30 nt; % of reads) in control and RA+BMP2-treated PGCs, split by antisense (blue) and sense (red) reads. Mean ± SD, n = 3 biological replicates, two-sided Welch t-test, n.s. B. Small RNA length distributions (18–40 nt) in control and RA+BMP2-treated PGCs, stacked by RNA biotype (miRNA, piRNA, tsRNA, rsRNA, sncRNA, mRNA, lncRNA, other). The piRNA-sized fraction (blue) was similar between conditions, indicating that meiotic induction does not detectably alter overall piRNA abundance (two-sided Welch t-test, n.s.). C. 5ʹ–5ʹ overlap-distance distributions for complementary piRNA pairs in treated (top) and control (bottom) PGCs. The y-axis shows the percentage of all overlapping pairs at each 5ʹ–5ʹ distance; the 10-nt overlap (red bar) is the ping-pong signature. A 10-nt peak is present in both conditions and its prominence does not differ (z = 19.6 ± 1.9 in control versus 19.2 ± 0.9 after treatment; two-sided Welch’s t-test, n.s.). Mean ± SD, n = 3 biological replicates.

**Table S1.** Differentially expressed genes between E10.5 (mitotic) and E16.5 (meiotic entry) chicken ovaries identified by DESeq2. Columns: Ensembl gene ID, base mean normalized count, log_2_ fold change, standard error, Wald statistic, p-value, Benjamini-Hochberg adjusted p-value, and direction of change (UP/DOWN). Genes with adjusted p < 0.05 and |log_2_FC| ≥ 1 are included. Three biological replicates per stage.

**Table S2.** Differentially expressed transposable element families between E10.5 and E16.5 ovaries, quantified using TEtranscripts with RepeatMasker annotation (galGal6). Columns: TE family name, base mean normalized count, log_2_ fold change, standard error, Wald statistic, p-value, Benjamini-Hochberg adjusted p-value, and functional category.

**Table S3.** Small-RNA and ping-pong metrics in zebrafish gonadal samples for zebrafish before meiosis I (3 wpf), at meiosis I (6 wpf), and in the mature ovary (24 wpf)

**Table S4.** Primer sequences used for qRT-PCR validation. Target genes include meiotic markers (*STRA8*, *SPO11*), piRNA pathway components (*PIWIL1*), the pluripotency marker *NANOG*, and reference genes (*GAPDH*, *ACTB*). Forward and reverse sequences, amplicon size, and annealing temperature are listed.

## Notes

### Competing Interest Statement

The authors have declared no competing interest.

### Summary of Updates

The support data, including Table S1-4, have been added to the manuscript, and a minor spelling problem has been fixed.

